# LeafContourEFD: a reproducible workflow for elliptic Fourier analysis with orientation normalization and lateral asymmetry

**DOI:** 10.64898/2026.04.16.718881

**Authors:** Kaede Konrai, Ryosuke Ito, Seiya Sunayama, Kurita Omura, Yuji Isagi, Kaoru Kitajima, Yusuke Onoda

## Abstract

**Premise:** Elliptic Fourier analysis is widely used to quantify leaf shape variation, but inconsistent normalization and orientation alignment can introduce biologically irrelevant variation. In addition, a reproducible workflow from raw images to normalized elliptic Fourier descriptors (EFDs) is still lacking.

**Methods and Results:** We developed LeafContourEFD, a GUI application for reproducible leaf morphometrics. It integrates image segmentation, contour extraction, EFD calculation, and an extended normalization framework, termed oriented true EFD normalization, based on a user-defined biological reference axis. Analyses of *Quercus serrata*, *Q. crispula*, and *Triadica sebifera* showed that existing normalization methods can introduce orientation-related variance when the first-harmonic major axis does not match the leaf base-to-tip axis. In contrast, oriented true normalization removed these artifacts, yielding clearer shape transitions along principal components allowing shape variation among leaves to be captured while preserving biologically meaningful lateral asymmetry.

**Conclusions:** LeafContourEFD improves interpretability and reproducibility in outline-based morphometrics and provides transparent outputs and metadata for data sharing and cross-study comparisons.

## INTRODUCTION

Leaf morphology exhibits remarkable variation not only among species but also within them (e.g., Ellis et al., 2009; Chitwood and Otoni, 2017; Fortini et al., 2025; Hightower et al., 2025). Such morphological diversity reflects a plant’s adaptation to its environment (Tsukaya, 2018; Yavas et al., 2024; Singhal et al., 2025), and has long been used to infer evolutionary strategies, taxonomy, and paleoclimate (Wolfe, 1978; Vogel, 2009; Wright et al., 2017; Wang et al., 2022). Recent advances in digital imaging and computational analysis have enabled quantitative assessment of leaf morphology from large amounts of image datasets, thereby accelerating research on the ecological and evolutionary significance of leaf shape (Chuanromanee et al., 2019; Gehan and Kellogg, 2017; Wilf et al., 2016). However, establishing an integrated workflow and ensuring reproducible quantification of morphometrics remain challenging (Chuanromanee et al., 2019; Powers and Hampton, 2019).

Geometric morphometric approaches have been widely applied to quantify leaf shape, and numerous software tools and programming packages have been developed to facilitate such analysis, including LeafJ (Maloof et al., 2013), LeafAnalyser (Weight et al., 2008), MASS (Chuanromanee et al., 2019), MORPHOJ (Klingenberg, 2011), SHAPE (Iwata, 2002), Momocs (Bonhomme et al., 2014), shapes (Dryden and Mardia, 2016), Geomorph (Adams and Otárola-Castillo, 2013), Tps Series (Rohlf, 2015), LeafMachine (Weaver et al., 2020), and LeafMachine2 (Weaver and Smith, 2023). However, most of these tools rely on pre-isolated leaf images or require extensive manual preprocessing, such as region-of-interest (ROI) extraction, rotation alignment, and contour cleaning. Consequently, a fully reproducible and automated pipeline extending from raw images to shape descriptors is still lacking.

Geometric morphometric methods can be broadly categorized into landmark-based and outline-based approaches (Kuhl and Giardina, 1982; Bookstein, 1992; Elewa, 2010; Budd, 2021). Landmark-based methods rely on homologous anatomical points (Bookstein, 1992), whereas outline-based approaches describe the entire contour without requiring predefined landmarks (Kuhl and Giardina, 1982; Dujardin et al., 2014; Hightower et al., 2025). Because leaf margins often lack clearly defined homologous points, outline-based methods are particularly suitable for quantifying leaf shape variation.

Among outline-based approaches, elliptic Fourier analysis (EFA) has become one of the most widely used techniques for quantifying biological outlines, as it captures the entire contour mathematically (Kuhl and Giardina, 1982; Jensen et al., 2002; Viscosi et al., 2009a, b; Mazo and Aribal, 2025). In EFA, normalization of elliptic Fourier descriptors (EFDs) or Fourier coefficients is essential to enable valid comparison of contour shapes among samples (Kuhl and Giardina, 1982; Ferson et al., 1985; Crampton, 1995; Bonhomme et al., 2014). This typically involves size standardization, orientation alignment, and starting-point correction (Ferson et al., 1985; Bonhomme et al., 2014; Wu et al., 2024). However, despite their well-established theoretical framework, consistent and reproducible implementation of these normalization procedures remains technically challenging.

Recently, Wu et al. (2024) proposed “true EFD normalization,” which mathematically resolves inconsistencies among previous EFD normalization methods. This approach adjusts the contour orientation by using the cross product of the first-order elliptic Fourier coefficients and corrects the x- and y-axis symmetries based on the second-order coefficients. In that study, Wu et al. (2024) also developed a graphical user interface (GUI) application, thereby establishing a complete workflow from image input to EFD extraction. However, this method still faces some challenges: (1) alignment may not be fully consistent when the major axes of the first-order ellipses differ among samples (i.e., when there is a mixture of leaves with aspect ratios lower and higher than 1); (2) biological orientations, such as the leaf base and tip, are not automatically standardized; and (3) there is limited ability to manually refine contour segmentation. While the software allows users to reconnect fragmented contours through raster-based editing and to apply basic morphological operations such as erosion and dilation, fine-grained control of individual boundary pixels for precise correction of segmentation artifacts is not fully supported.

In this study, we developed a GUI application built on napari (Sofroniew et al., 2025) that provides a fully reproducible workflow for leaf morphometrics. The application integrates all steps—from segmentation and contour extraction to EFD calculation and an extended true EFD normalization framework based on the work of Wu et al. (2024)—within a single interactive environment. It allows both conventional and true EFDs to be obtained simultaneously while aligning all samples according to a user-defined biological orientation (e.g., leaf base–tip direction). It also incorporates deep-learning-based segmentation using the Segment Anything Model 2 (SAM2; Ravi et al., 2024) and automatically records complete metadata (e.g., contour, masks both original and hand-edited, landmarks, scale, and cropped coordinates from the original image) to ensure transparency and reproducibility (Powers and Hampton, 2019). By combining a rich interactive GUI with mathematically rigorous shape normalization (Selzer et al., 2023; Wu et al., 2024), this open-source tool provides a user-friendly and extensible platform for reproducible morphometric analysis of leaves and other biological structures.

## METHODS AND RESULTS

### Software overview

LeafContourEFD is an open-source program (BSD 3-Clause License) written in Python 3 and built on napari (Sofroniew et al., 2025). The source code, user manual, and example data set are freely available at https://github.com/maple60/leaf-contour-efd. The software is available for Windows, macOS, and Linux, either as a stand-alone executable build with PyInstaller 6.16.0 or from the source code. If users wish to apply the SAM2 (Ravi et al., 2024) for contour-area segmentation, the installation instructions and script files for SAM2 are also available in the repository.

A schematic overview of the analytical workflow—from image loading, ROI selection, landmark placement, segmentation, and contour extraction to EFD calculations—is presented in Figure 1. This figure outlines the major processing steps and their corresponding modules implemented in LeafContourEFD.

**Figure 1.**
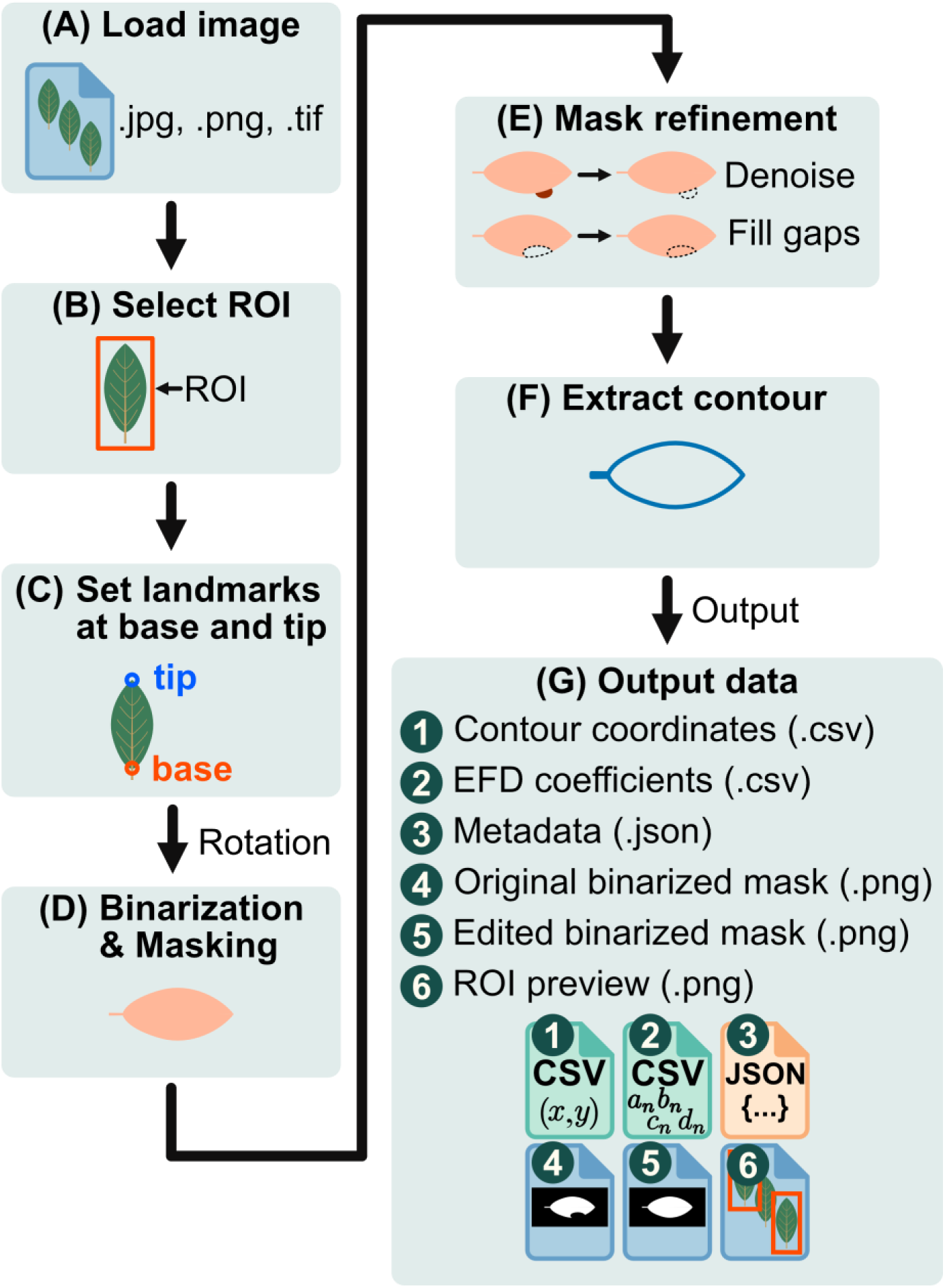
Workflow of LeafContourEFD, showing the sequential procedures from image acquisition to elliptic Fourier descriptor (EFD) computation and metadata export.

### Input preparation

The minimum input required for LeafContourEFD is a leaf image in a standard image format (e.g., PNG, JPEG, or TIFF) (Figure 1A). Although the program does not strictly require a specific image background, images with a plain white background—such as scanned specimens—are recommended for accurate and efficient segmentation and subsequent contour extraction. For scanned leaves, a resolution of 300 dpi (dots per inch) is generally sufficient for morphometric analysis.

When a plain background is available, segmentation can be performed using the built-in thresholding method of Otsu (1979) followed by optional manual threshold adjustment. For images with more complex or textured background, SAM2 provides a more robust solution for regional segmentation.

When the analysis involves leaf-area measurement, users can apply scale calibration. This can be achieved by including a visible scale bar in the image or by specifying the DPI value of the scanned image.

### Core functions

#### Region-of-interest selection and cropping

To exclude irregular or unsuitable contours in subsequent analysis (e.g., large missing margins or abnormal bending), LeafContourEFD adopts an ROI-based selection step prior to contour extraction (Figure 1B). By defining an ROI that encloses the target object, users can visually confirm whether the specimen is appropriate for contour analysis before proceeding. This step helps remove objects that could otherwise produce erroneous contours in fully automated processing.

During ROI selection, small extraneous objects such as dust within the ROI do not affect the analysis because the program automatically extracts only the largest closed contour within the selected ROI. Minor defects or noise—such as small missing parts along the leaf edge, attached dust particles, or frayed fibers from the end of a petiole—can be manually corrected after generation of the binary mask, if necessary.

Once the ROI is confirmed, the selected region is automatically cropped and passed to the landmark placement and rotation module for subsequent processing.

#### Landmark placement and image rotation

On the cropped ROI image, two landmarks are placed at the base and tip of the leaf blade (or the target object) (Figure 1C). Then, the image is automatically rotated so that the base is positioned on the left, the tip on the right, and the line connecting them becomes horizontal (Figure 1D). This process allows all samples to be aligned according to their primary axis, facilitating shape comparison among specimens. The rotation step corresponds to the rotation normalization procedure in previous EFA-based methods (Kuhl and Giardina, 1982; Wu et al., 2024).

#### Binarization and segmentation

To define the region from which contours are extracted, LeafContourEFD generates a binarized mask for segmentation (Figure 1D). Two alternative methods are available for mask generation: Otsu’s thresholding method (Otsu, 1979) and SAM2 (Ravi et al., 2024). Otsu’s method is much faster than SAM2 and performs particularly well when the image background is plain white (e.g., scanned image, Figure 1A). After automatic binarization, users can manually adjust the threshold value to refine the segmentation if needed (Figure 1E). SAM2 is more robust when the image background is non-white or when multiple objects overlap within the field of view. Please note that SAM2 can work well even when Otsu’s method performs well; however, it takes longer and occasionally omits serrated leaf margins or small teeth.

After generating the binarized mask, users can refine the outline using the built-in editing tools provided by napari (Sofroniew et al., 2025; Figure 1E). If noise overlaps the contour, it can be removed with the eraser tool. When users wish to perform contour comparisons excluding the petiole, the eraser tool can also be used to separate the leaf blade from the petiole. In cases where small gaps occur along the leaf margin or the contour is not completely closed, the brush and polygon tools allow users to manually reconstruct or complete the outline.

#### Contour extraction

Based on the generated binarized mask, the x-y coordinates of the contour are extracted (Figure 1F). Because the extracted contour is displayed together with the mask in the graphical user interface (GUI), users can visually inspect the result. If extraction errors or artifacts are detected, users can delete the contour, generate the mask, and extract the contour again (Figure 1E, F).

Contour extraction begins at the point with the maximum x-axis value, which corresponds to the tip of the leaf blade. This ensures that all samples share a biologically consistent starting point for shape analysis, resulting in more robust comparisons among specimens. This step corresponds to the starting point normalization procedure in previous EFA-based methods (Kuhl and Giardina, 1982).

#### Elliptic Fourier descriptor calculation

From the extracted contour, both the EFDs and the biological primary axis-oriented true normalized EFDs are automatically calculated and exported together with the contour coordinates as CSV files. Because the original contour coordinate data are also exported, users can independently compute EFDs or perform additional preprocessing before further analysis.

The exported data are compatible with the R package Momocs (Bonhomme et al., 2014) for EFA, enabling seamless integration with downstream statistical analyses such as principal component analysis (PCA). The output files and example analysis are provided in the public repository associated with the present study.

The EFDs are computed following the standard procedure described by Kuhl and Giardina (1982). Size, rotation, and x-axis reflection —which is essential for avoiding sign reversals caused by mirror-reflected contours— are normalized according to the method of Wu et al. (2024). Unlike conventional implementations that rely on the orientation of the first harmonic ellipse, LeafContourEFD standardizes the rotation and starting point based on biological primary axis predefined landmarks, ensuring consistent alignment among samples without the need for harmonic-based normalization.

#### Results and metadata export

In addition to the contour coordinates and EFDs, LeafContourEFD exports metadata and intermediate images to ensure the reproducibility and transparency of the analysis (Powers and Hampton, 2019; Figure 1G).

The exported metadata includes the path of the original image, coordinates of the cropped ROI, landmark positions, rotation angle, binarization method, and the threshold value (if Otsu’s method is selected). The intermediate images consist of the cropped ROI, rotated image, automatically generated mask, and manually edited mask.

### Example workflow

#### Species and image acquisition

We selected three tree species to demonstrate the utility of our pipeline: *Quercus serrata*, *Q. crispula*, and *Triadica sebifera* (Appendix S1; see Supporting Information with this article). A total of 462 leaves from 102 individuals of *Q. serrata*, 284 leaves from 92 individuals of *Q. crispula*, and 30 leaves from 2 individuals of *T. sebifera* were analyzed. Leaves of *Q. serrata* and *Q. crispula* were collected from Mt. Jatanigamine, Shiga, Japan (35.3204 °N, 135.9286 °E), whereas *T. sebifera* leaves were collected from the campus of Kyoto University, Kyoto, Japan (35.0257 °N, 135.7792 °E).

*Quercus serrata* and *Q. crispula* are closely related species that can form natural hybrids and share similar overall leaf morphology, with elongated, toothed leaves (Ohba, 1989). However, Q. serrata typically has a longer petiole (ca. 5-20 mm) and its maximum leaf with lies near the center of the lamina, whereas *Q. crispula* has a shorter petiole (ca. 1-5mm) and its maximum leaf width is positioned more distally, around the distal two-third of the lamina. These two *Quercus* species were thus used to evaluate how effectively leaf shape differences can be visualized by EFDs and subsequent PCA.

In contrast, *T. sebifera* often bears broad, nearly orbicular leaves, with leaf with frequently exceeding leaf length measured along the midrib. This species was therefore included to assess the robustness of our rotation normalization procedure. Because *T. sebifera* has a long petiole, we removed the petiole region using our software before the morphometric analysis.

All leaves were scanned using a flatbed scanner (GT-S650 or GT-X970, Epson) at 300 dpi on a white background to obtain high-contrast digital images suitable for contour extraction

#### Outline extraction and orientation normalization

After contour extraction using LeafContourEFD, each contour was randomly rotated from 0° to 360° to evaluate the robustness of the normalization procedure. Elliptic Fourier descriptors were then computed using the efourier function of the Momocs package (Bonhomme et al., 2014) with norm = TRUE in R 4.5.3 (R Core Team, 2025). True EFD normalization was subsequently applied following the procedure of Wu et al. (2024), including both x- and y-axis symmetric normalization. Reconstructed contours were generated from the normalized coefficients using the efourier_shape function of Momocs. The number of harmonics used for reconstruction was set to 35 for *Q. serrata* and *Q. crispula* and 10 for *T. sebifera*, reflecting differences in leaf-margin complexity:the *Quercus* species have serrated margins, whereas *T. sebifera* has entire margins.

In addition, we implemented a procedure that further aligns the reconstructed contours to the biological primary axis of each leaf, using the landmark-defined axis as the directional reference. Hereafter, we refer to this procedure as oriented true EFD normalization. The normalized coefficients generated by this procedure are automatically exported from LeafContourEFD for subsequent analyses.

All contours reconstructed from oriented true EFD normalization coefficients were aligned in the same direction as the original contours extracted by LeafContourEFD (Figure 2A, Figure 3A). Consistency is critical here because inconsistent orientation introduces biologically irrelevant variation, which may distort multivariate analysis such as PCA and bias average shape reconstruction.

**Figure 2.**
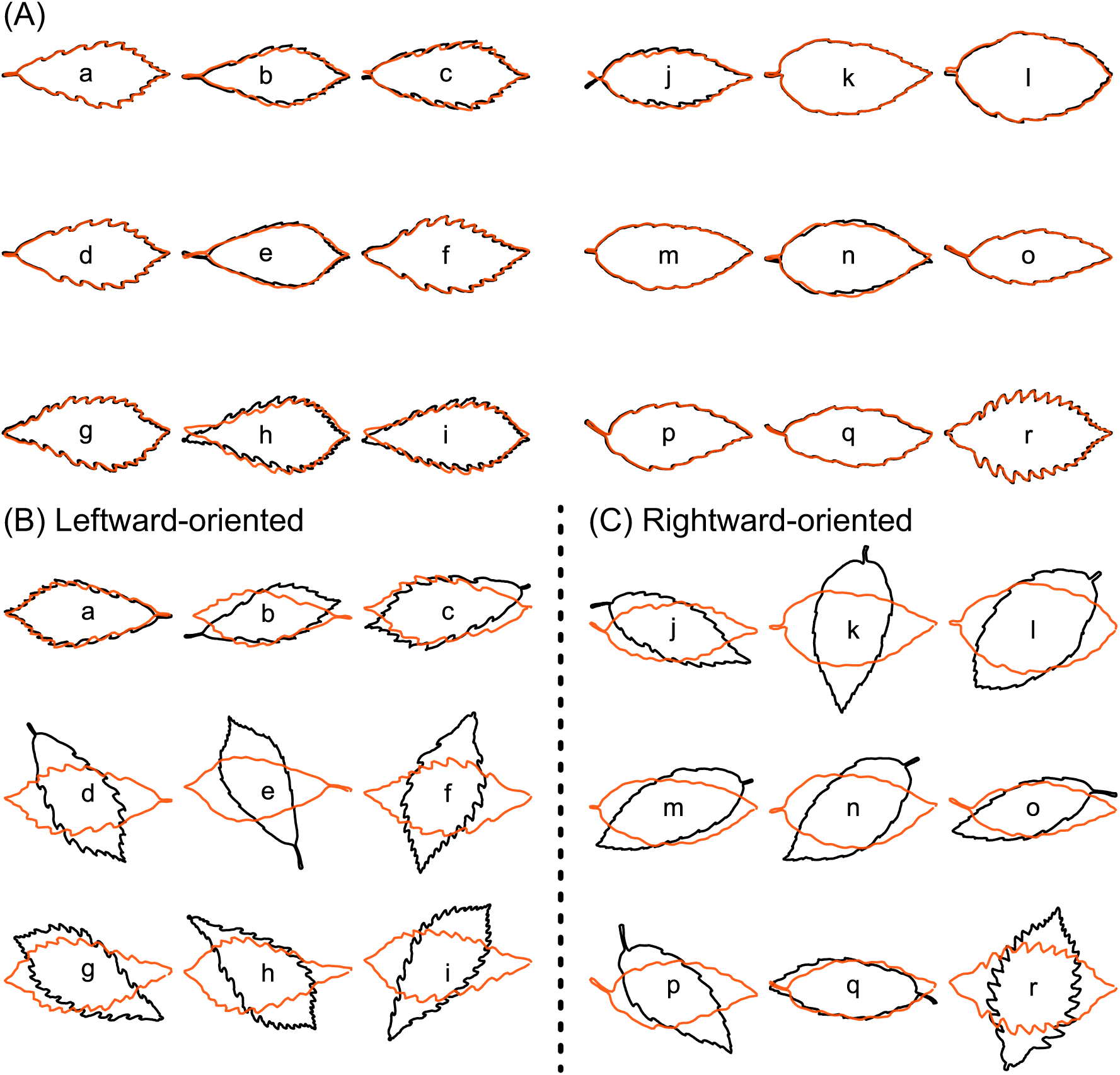
Comparison between the extracted contours and those reconstructed from normalized EFD coefficients of *Quercus serrata* and *Q. crispula*. Black lines represent the original contours and orange lines indicate the reconstructed ones. (A) Contours exported from LeafContourEFD and those reconstructed using the oriented true EFD normalization. (B, C) Randomly rotated contours and those reconstructed from the coefficients obtained by the true EFD normalization (Wu et al., 2024). (B) Leftward and (C) rightward oriented reconstructions. Lower case letters inside the contours correspond to the same samples across panels, allowing direct comparison between the oriented and conventional normalizations. Labels a–e and j–q represent *Q. serrata*, whereas f–i and r represent *Q. crispula*.

**Figure 3.**
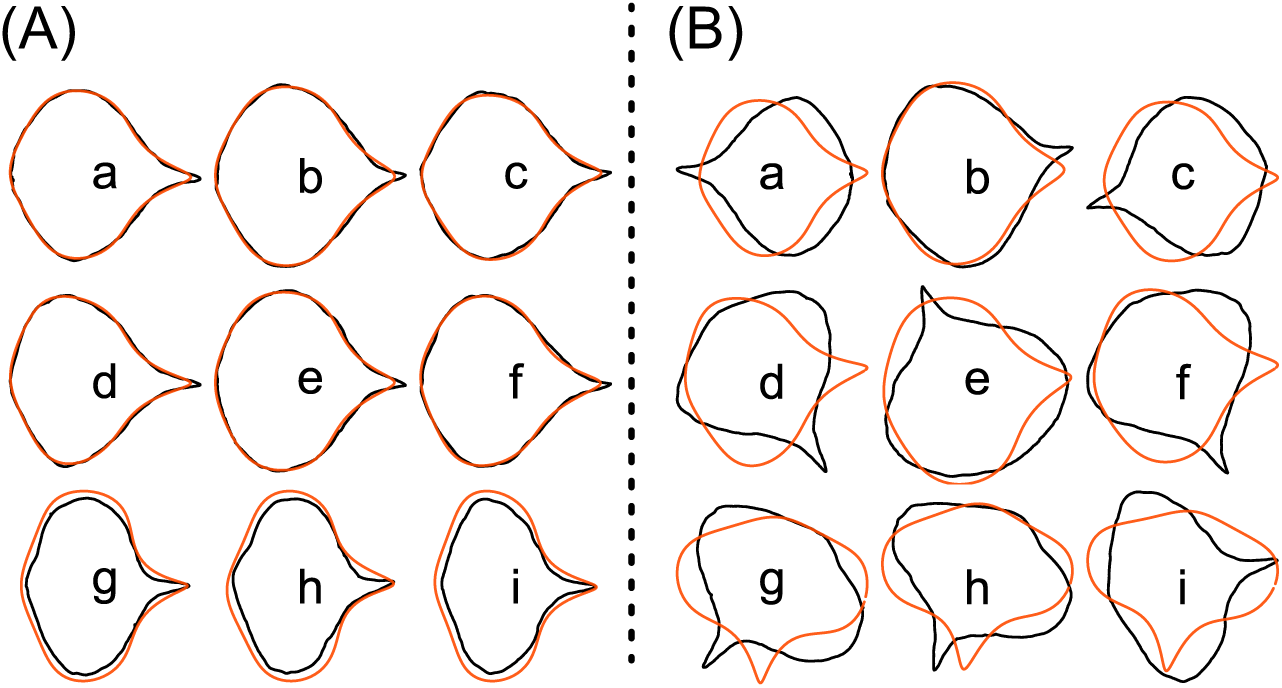
Comparison of reconstructed leaf contours of *Triadica sebifera* between different normalization methods. Black lines represent the original contours, and orange lines represent the reconstructed ones. (A) Original and reconstructed contours obtained using the oriented true EFD normalization implemented in LeafContourEFD. (B) Original and reconstructed contours obtained using the true EFD normalization following Wu et al. (2024). Lower case panel labels (a–i) indicate corresponding samples between (A) and (B).

In contrast, under the conventional true EFD normalization, leftward- and rightward-oriented leaves were occasionally mixed. For example, among 462 *Q. serrata* samples, eight leaves were oriented to the right, while the others were oriented to the left. Similarly, among 284 *Q. crispula* samples, one leaf was oriented to the right while the others were to the left (Figure 2B, C). For *T. sebifera*, 9 of 30 contours were reconstructed vertically or obliquely relative to the leaf blade (Figure 3B).

In all cases, our oriented true EFD normalization successfully unified the contour orientation with the original direction of each leaf, achieving a consistent biological orientation across samples.

#### Principal component analysis of EFDs

We performed PCA based on three sets of normalized coefficients: (1) conventional EFD normalization (Kuhl and Giardina, 1982; Bonhomme et al., 2014), (2) true EFD normalization (Wu et al., 2024), and (3) oriented true EFD normalization implemented in LeafContourEFD. In the software, each contour is already normalized with respect to direction and starting point prior to analysis. To evaluate the robustness of the normalization procedures used in the conventional and true EFD methods, we artificially rotated each outline by a random angle between 0° and 360° and randomly shifted the starting point prior to applying these two normalization approaches. PCA was conducted using the PCA function of the Momocs package (Bonhomme et al., 2014), and contour variations along principal axes (PC1 and PC2) were visualized using plot_PCA.

For *Q. serrata* and *Q. crispula*, all three normalization procedures separated the two species along PC1 (Figure 4A–C). Contours on the negative side of PC1 showed an ovate outline with a longer petiole (*Q. serrata*-like), whereas those on the positive side of PC1 exhibited a broader upper blade with an indistinct petiole (*Q. crispula*-like). The proportions of variance explained by PC1 and PC2 were 43.3% and 23.1% for the conventional normalization, 55.2% and 10.6% for the true EFD normalization, and 44.7% and 14.8% for the oriented true EFD normalization, respectively. The higher explanatory power of PC1 in the true EFD normalization than in the oriented true EFD normalization likely reflects the fact that, in the former, PC1 captured variation associated with the major axis of the first harmonic ellipse, regardless of whether this axis matched the biologically correct leaf orientation. Actual leaves, meanwhile, can be broader than long; thus, their major axis did not correspond to the major axis of the first harmonic ellipse, which can result in a low PC1 score in the oriented true EFD normalization. Similarly, the higher explanatory power of PC2 in the conventional normalization than in the oriented true EFD normalization was because the second axis of the conventional normalization captured variance associated with left–right asymmetry, which was normalized before the analysis in the oriented true EFD normalization. In short, the oriented true EFD normalization can provide more biologically meaningful results.

**Figure 4.**
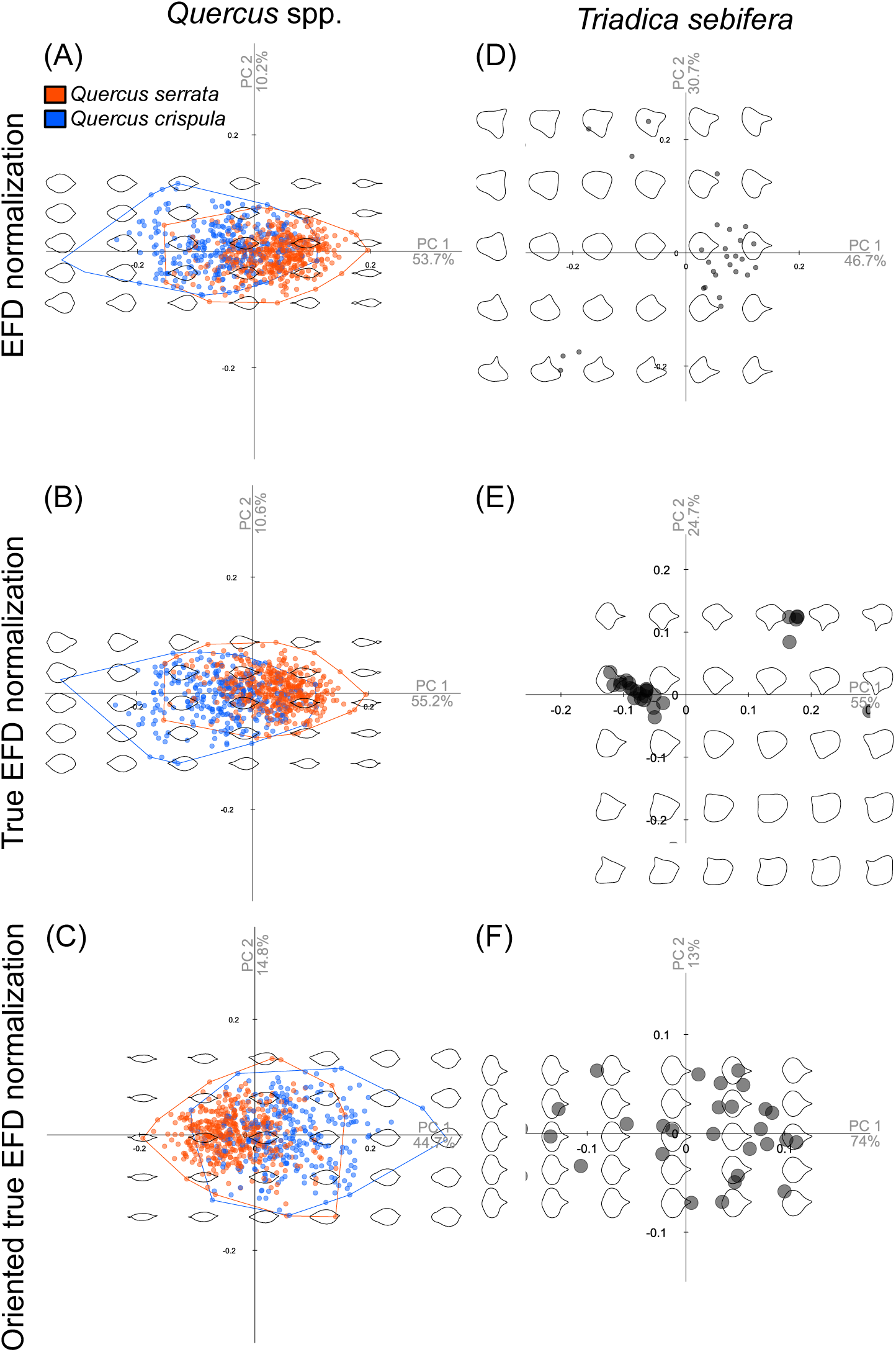
Results of principal component analysis (PCA) based on normalized EFD coefficients obtained by different normalization methods. (A, D) Conventional normalization following Kuhl and Giardina (1982), (B, E) true EFD normalization following Wu et al. (2024), and (C, F) oriented true EFD normalization implemented in LeafContourEFD. Each point represents a single leaf contour, and polygons in panels (A–C) indicate convex hulls encompassing the samples of each species. Background outlines in each plot show contour reconstructions along the principal component (PC) axes. Panels A–C correspond to *Quercus serrata* (orange) and *Q. crispula* (blue), whereas panels D–F show the results for *Triadica sebifera*. Percent variance explained is shown below the PC1 and PC2 axes.

This difference became more pronounced in *T. sebifera*, whose leaves are often broader than they are long (Appendix S1). Under conventional and true EFD normalization, mismatches between the geometric major axis and the midrib direction generated orientation-related outliers, causing PC1 to capture variance associated with leaf orientation rather than intrinsic shape differences (Figure 4D, E). In contrast, the oriented true EFD normalization yielded a clear transition from broad to elongated leaf shapes, explaining 74.0% of total variance compared with substantially lower values (46.7% in the conventional method and 55.0% in true EFD normalization) with other methods (Figure 4F). Together, these results demonstrate that the oriented true EFD reduced orientation-induced variance and provided more biologically interpretable shape differentiation.

## CONCLUSIONS

Leaf shape provides information not only for species identification but also for understanding relationships between leaf shape and functions, making it a physiologically, ecologically and evolutionarily important characteristic (Li et al., 2018; Wang et al., 2022; Kassout et al., 2024). With recent advances in imaging technology and the digitization of herbarium collections, large numbers of leaf images can be obtained without difficulty. However, extracting quantitative and comparable information on leaf outlines from digital images has been challenging due to issues with preprocessing, reproducibility, and analytical standardization (Ferson et al., 1985; Powers and Hampton, 2019; Wu et al., 2024). LeafContourEFD can provide solutions to these challenges by implementing a user-friendly GUI and oriented true EFD normalization. This method enables robust and reproducible shape comparison across diverse leaf morphologies. In addition, the software was designed to ensure user-friendliness and reproducibility, providing transparent metadata and intermediate outputs that facilitate data sharing, comparison, and reanalysis across studies. We expect that LeafContourEFD will become a practical and reproducible platform for digital leaf morphometrics, promoting the development of databases and advancing the integration of leaf shape information into evolutionary and ecological research.

## ACKNOWLEDGEMENTS

This work was supported by JSPS KAKENHI (Grant Number 21H05314) and JST SPRING (Grant Number JPMJSP2110). We thank Tom Buckle from Scribendi (www.scribendi.com) for editing a draft of this manuscript.

## AUTHOR CONTRIBUTIONS

K. Konrai conducted the analyses and wrote the initial manuscript. R.I., S.S., K.O, Y.I., and Y.O. collected leaf samples in the field. K. Kitajima supervised the overall research design and contributed to project planning. Y.O. supervised the study and provided critical revisions to the manuscript. All authors approved the final version of the manuscript.

## DATA AVAILABILITY

All contour coordinate data and normalized EFD coefficients used for statistical analyses are available in the Dryad Digital Repository at https://doi.org/10.5061/dryad.6t1g1jxdh. The source code of LeafContourEFD is available on GitHub (https://github.com/maple60/leaf-contour-efd) and archived on Zenodo (https://doi.org/10.5281/zenodo.19449129). The analysis code is available on GitHub (https://github.com/maple60/leaf-contour-efd-analysis) and archived on Zenodo (https://doi.org/10.5281/zenodo.19490591).

## SUPPORTING INFORMATION

Additional Supporting Information may be found online in the Supporting Information section at the end of the article.

Appendix S1. Scanned leaf images of the target species: (A) *Quercus serrata*, (B) *Quercus crispula*, and (C) *Triadica sebifera*.

## REFERENCES

1. Adams, D. C., and Otárola-Castillo, E. 2013. geomorph: An R package for the collection and analysis of geometric morphometric shape data. Methods in Ecology and Evolution 4(4), 393–399. 10.1111/2041-210X.12035

2. Bonhomme, V., Picq, S., Gaucherel, C., and Claude, J. 2014. Momocs: Outline analysis using *R*. Journal of Statistical Software 56(13), 1–24. 10.18637/jss.v056.i13

3. Bookstein, F. L. 1992. Morphometric tools for landmark data: Geometry and biology. Cambridge University Press, Cambridge, UK. 10.1017/CBO9780511573064

4. Budd, G. E. 2021. Morphospace. Current Biology 31(19), R1181–R1185. 10.1016/j.cub.2021.08.040

5. Chitwood, D. H., and Otoni, W. C. 2017. Divergent leaf shapes among *Passiflora* species arise from a shared juvenile morphology. Plant Direct 1(5), e00028. 10.1002/pld3.28

6. Chuanromanee, T. S., Cohen, J. I., and Ryan, G. L. 2019. Morphological Analysis of Size and Shape (MASS): An integrative software program for morphometric analyses of leaves. Applications in Plant Sciences 7(9), e11288. 10.1002/aps3.11288

7. Crampton, J. S. 1995. Elliptic Fourier shape analysis of fossil bivalves: Some practical considerations. Lethaia 28(2), 179–186. 10.1111/j.1502-3931.1995.tb01611.x

8. Dryden, I. L., and Mardia, K. V. 2016. Statistical shape analysis with applications in R (2nd edition). John Wiley and Sons, Hoboken, NJ, USA.

9. Dujardin, J.-P., Kaba, D., Solano, P., Dupraz, M., McCoy, K. D., and Jaramillo-O, N. 2014. Outline-based morphometrics, an overlooked method in arthropod studies? Infection, Genetics and Evolution 28, 704–714. 10.1016/j.meegid.2014.07.035

10. Elewa, A. M. T. (Ed.). 2010. Morphometrics for nonmorphometricians (Vol. 124). Springer, London, UK. 10.1007/978-3-540-95853-6

11. Ellis, B., Daly, D., Hickey, L., Johnson, K., Mitchell, J., Wilf, P., and Wing, S. 2009. Manual of leaf architecture. Cornell University Press CABI, Ithaca, NY, USA. 10.1079/9781845935849.0000

12. Ferson, S., Rohlf, F. J., and Koehn, R. K. 1985. Measuring shape variation of two-dimensional outlines. Systematic Zoology 34(1), 59–68. 10.2307/2413345

13. Fortini, P., Proietti, E., Stojnic, S., Di Marzio, P., Aravanopoulos, F. A., Benavides, R., Loy, A., and Di Pietro, R. 2025. First results of a geometric morphometric analysis of the leaf size and shape variation in *Quercus petraea* across a wide European area. Forests 16(1), 70. 10.3390/f16010070

14. Gehan, M. A., and Kellogg, E. A. 2017. High-throughput phenotyping. American Journal of Botany 104(4), 505–508. 10.3732/ajb.1700044

15. Hightower, A., Hall, S., Camacho, R. U., Papamichail, A., Adamski, E., Colligan, C., Deneen, A., et al. 2025. Procrustean pseudo-landmark methods in Python to measure massive quantities of leaf shape data (p. 2025.08.08.669192). bioRxiv. 10.1101/2025.08.08.669192

16. Iwata, H. 2002. SHAPE: A computer program package for quantitative evaluation of biological shapes based on elliptic Fourier descriptors. Journal of Heredity 93(5), 384–385. 10.1093/jhered/93.5.384

17. Jensen, R. J., Ciofani, K. M., and Miramontes, L. C. 2002. Lines, outlines, and landmarks: Morphometric analyses of leaves of *Acer rubrum*, *Acer saccharinum* (Aceraceae) and their hybrid. Taxon 51(3), 475–492. 10.2307/1555066

18. Kassout, J., Terral, J.-F., Souali, H., and Ater, M. 2024. Environment-dependent and intraspecific variations in leaf and size traits of a native wild olive (*Olea europaea* L.) along an aridity gradient in Morocco: A functional perspective. Plant Ecology 225(9), 943–959. 10.1007/s11258-024-01445-2

19. Klingenberg, C. P. 2011. MORPHO J: An integrated software package for geometric morphometrics. Molecular Ecology Resources 11(2), 353–357. 10.1111/j.1755-0998.2010.02924.x

20. Kuhl, F. P., and Giardina, C. R. 1982. Elliptic Fourier features of a closed contour. Computer Graphics and Image Processing 18(3), 236–258. 10.1016/0146-664X(82)90034-X

21. Li, M., An, H., Angelovici, R., Bagaza, C., Batushansky, A., Clark, L., Coneva, V., et al. 2018. Topological data analysis as a morphometric method: Using persistent homology to demarcate a leaf morphospace. Frontiers in Plant Science 9, 553. 10.3389/fpls.2018.00553

22. Maloof, J. N., Nozue, K., Mumbach, M. R., and Palmer, C. M. 2013. LeafJ: An ImageJ plugin for semi-automated leaf shape measurement. Journal of Visualized Experiments 71, e50028. 10.3791/50028

23. Mazo, K. R. F., and Aribal, L. G. 2025. Leaf size indices and outline-based geomorphometric analysis of five Philippine endemic *Saurauia* Willd. (Actinidiaceae). Jurnal Sylva Lestari 13(2), Article 2. 10.23960/jsl.v13i2.1139

24. Ohba, H. 1989. New names and notes of Japanese woody plants. The Journal of Japanese Botany 64(11), 321–329. 10.51033/jjapbot.64_11_8388

25. Otsu, N. 1979. A threshold selection method from gray-level histograms. *IEEE Transactions on Systems*, Man, and Cybernetics 9(1), 62–66. 10.1109/TSMC.1979.4310076

26. Powers, S. M., and Hampton, S. E. 2019. Open science, reproducibility, and transparency in ecology. Ecological Applications 29(1), e01822. 10.1002/eap.1822

27. R Core Team. 2025. R: A language and environment for statistical computing. R Foundation for Statistical Computing. https://www.R-project.org/

28. Ravi, N., Gabeur, V., Hu, Y.-T., Hu, R., Ryali, C., Ma, T., Khedr, H., et al. (2024). SAM 2: Segment anything in images and videos (arXiv:2408.00714). arXiv. 10.48550/arXiv.2408.00714

29. Rohlf, F. J. 2015. The tps series of software. Hystrix, the Italian Journal of Mammalogy 26(1), 9–12. 10.4404/hystrix-26.1-11264

30. Selzer, G. J., Rueden, C. T., Hiner, M. C., Evans, E. L., Harrington, K. I. S., and Eliceiri, K. W. 2023. napari-imagej: ImageJ ecosystem access from napari. Nature Methods 20(10), 1443–1444. 10.1038/s41592-023-01990-0

31. Singhal, Y. K., Boyle, J. A., and Stinchcombe, J. R. 2025. Differences in stomatal conductance between leaf shape genotypes of *Ipomoea hederacea* suggest divergent ecophysiological strategies. microPublication Biology. 10.17912/micropub.biology.001528

32. Sofroniew, N., Lambert, T., Bokota, G., Nunez-Iglesias, J., Sobolewski, P., Sweet, A., Gaifas, L., et al. 2025. napari: A multi-dimensional image viewer for Python (Version v0.6.6) [Computer software]. Zenodo. 10.5281/ZENODO.17367124

33. Tsukaya, H. 2018. Leaf shape diversity with an emphasis on leaf contour variation, developmental background, and adaptation. *Seminars in Cell and Developmental Biology*, Shape and Form in Plant Development 79, 48–57. 10.1016/j.semcdb.2017.11.035

34. Viscosi, V., Fortini, P., Slice, D. E., Loy, A., and Blasi, C. 2009a. Geometric morphometric analyses of leaf variation in four oak species of the subgenus *Quercus* (Fagaceae). Plant Biosystems 143(3), 575–587. 10.1080/11263500902775277

35. Viscosi, V., Lepais, O., Gerber, S., and Fortini, P. 2009b. Leaf morphological analyses in four European oak species (*Quercus*) and their hybrids: A comparison of traditional and geometric morphometric methods. Plant Biosystems 143(3), 564–574. 10.1080/11263500902723129

36. Vogel, S. 2009. Leaves in the lowest and highest winds: Temperature, force and shape. New Phytologist 183(1), 13–26. 10.1111/j.1469-8137.2009.02854.x

37. Wang, H., Wang, R., Harrison, S. P., and Prentice, I. C. 2022. Leaf morphological traits as adaptations to multiple climate gradients. Journal of Ecology 110(6), 1344–1355. 10.1111/1365-2745.13873

38. Weaver, W. N., Ng, J., and Laport, R. G. 2020. LeafMachine: Using machine learning to automate leaf trait extraction from digitized herbarium specimens. Applications in Plant Sciences 8(6), e11367. 10.1002/aps3.11367

39. Weaver, W. N., and Smith, S. A. 2023. From leaves to labels: Building modular machine learning networks for rapid herbarium specimen analysis with LeafMachine2. Applications in Plant Sciences 11(5), e11548. 10.1002/aps3.11548

40. Weight, C., Parnham, D., and Waites, R. 2008. Technical advance: LeafAnalyser: A computational method for rapid and large-scale analyses of leaf shape variation. The Plant Journal 53(3), 578–586. 10.1111/j.1365-313X.2007.03330.x

41. Wilf, P., Zhang, S., Chikkerur, S., Little, S. A., Wing, S. L., and Serre, T. 2016. Computer vision cracks the leaf code. Proceedings of the National Academy of Sciences 113(12), 3305–3310. 10.1073/pnas.1524473113

42. Wolfe, J. A. 1978. A paleobotanical interpretation of tertiary climates in the Northern Hemisphere: Data from fossil plants make it possible to reconstruct Tertiary climatic changes, which may be correlated with changes in the inclination of the earth’s rotational axis. American Scientist 66(6), 694–703.

43. Wright, I. J., Dong, N., Maire, V., Prentice, I. C., Westoby, M., Díaz, S., Gallagher, R. V., et al. 2017. Global climatic drivers of leaf size. Science 357(6354), 917–921. 10.1126/science.aal4760

44. Wu, H., Yang, J.-J., Li, C.-Q., Ran, J.-H., Peng, R.-H., and Wang, X.-Q. 2024. Reliable and superior elliptic Fourier descriptor normalization and its application software ElliShape with efficient image processing (arXiv:2412.10795). arXiv. 10.48550/arXiv.2412.10795

45. Yavas, I., Jamal, M. A., Din, K. U., Ali, S., Hussain, S., and Farooq, M. 2024. Drought-induced changes in leaf morphology and anatomy: Overview, implications and perspectives. Polish Journal of Environmental Studies 33(2), 1517–1530. 10.15244/pjoes/174476

