## Supporting information for "LeafContourEFD: a reproducible workflow for elliptic Fourier analysis with orientation normalization and lateral asymmetry"

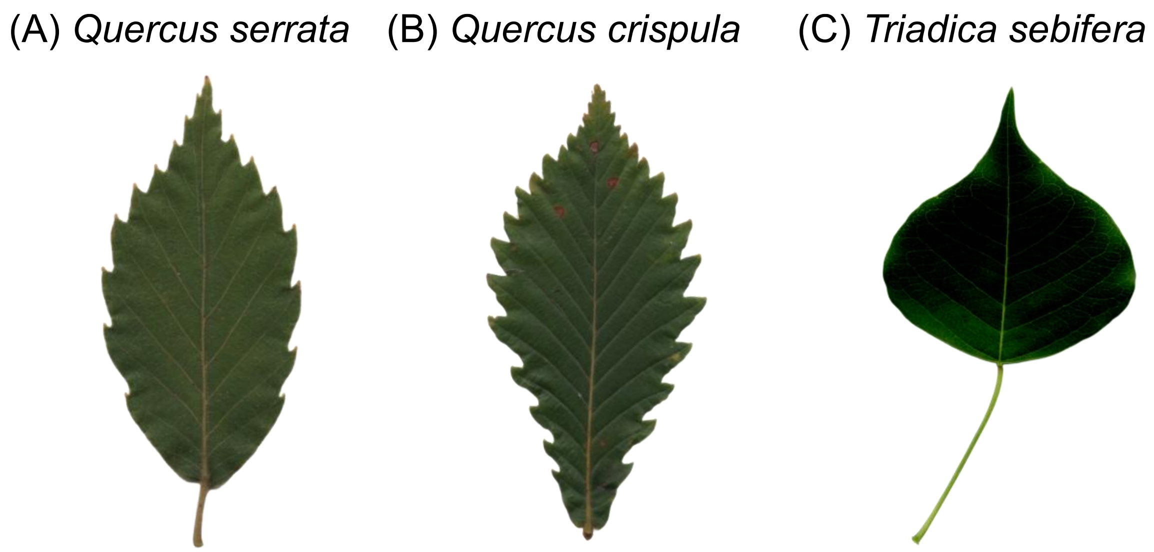


**Figure S1.** Scanned leaf images of the target species: (A) *Quercus serrata*, (B) *Quercus crispula*, and (C) *Triadica sebifera*.
